# Integrating multiple single-cell multi-omics samples with Smmit

**DOI:** 10.1101/2023.04.06.535857

**Authors:** Changxin Wan, Zhicheng Ji

## Abstract

Multi-sample single-cell multi-omics datasets, which simultaneously measure multiple data modalities in the same cells across multiple samples, facilitate the study of gene expression, gene regulatory activities, and protein abundances on a population scale. We developed Smmit, a computational method for integrating data both across samples and modalities. Compared to existing methods, Smmit more effectively removes batch effects while preserving relevant biological information, resulting in superior integration outcomes. Additionally, Smmit is more computationally efficient and builds upon existing computational pipelines, requiring minimal effort for implementation. Smmit is an R software package that is freely available on Github: https://github.com/zji90/Smmit.

## Introduction

Single-cell multi-omics sequencing measures multiple modalities of molecular profiles in the same cells. Examples of such technologies include joint profiling of gene expression and protein abundances (e.g., CITE-seq^1^) and joint profiling of gene expression and chromatin accessibility (e.g., 10x Multiome, SHARE-seq^2^ and SNARE-seq^3^). Multi-sample single-cell multiomics datasets have been generated to study and compare gene expression and gene regulatory activities across samples in neural development^4^, leukemia^5^, skin fibroblast^6^, and other biological systems. These datasets enable population-level studies of cells’ comprehensive functions and characteristics under different conditions, providing more in-depth insights than data derived from a single modality or a single sample.

To effectively analyze single-cell sequencing data from multiple samples and modalities, ideally, one needs to first integrate information across samples and modalities to generate a single representation of reduced dimensions. The integration process harmonizes discrepancies across samples and modalities, facilitating downstream analysis such as cell clustering, cell type identification, and characterization of regulatory behavior within cell subpopulations. Several integration methods have been developed for this purpose. Multigrate^7^ uses a generative multi-view neural network to learn a joint latent space from multiple modalities while accounting for technical biases within each modality. scVAEIT^8^ uses a probabilistic variational autoencoder model to integrate and impute multimodal datasets with mosaic measurements. scMoMaT^9^ uses matrix tri-factorization to integrate single cell multi-omics data under the mosaic integration scenario. MultiVI^10^ uses a deep generative model for probabilistic analysis and integration of multimodal datasets. MOFA+^11^ uses a statistical framework based on variational inference for reconstructing an integrated low-dimensional representation of the single-cell multi-modal data. totalVI^12^ uses a deep generative model that enables integration and multifaceted analysis of CITE-seq data.

A major limitation of these methods is their computational inefficiency. Most of these approaches rely on complex deep neural networks and statistical modeling. As the cost of single-cell sequencing continues to decrease and dataset sizes continue to grow, these methods may become computationally impractical due to the extensive time and memory required to complete the computations. Moreover, these methods often necessitate dedicated computational resources and advanced computational expertise for implementation. For instance, methods based on deep neural networks require access to GPUs and proficiency in platforms such as PyTorch. These requirements hinder the democratization of complex single-cell multi-omics data analysis, particularly in research groups lacking sufficient computational resources.

To address this challenge, we developed Smmit, a computationally efficient method for <u>s</u>ingle-cell <u>mu</u>lti-sample and <u>mu</u>lti-omics <u>i</u>n<u>t</u>egration (Methods). Smmit is a two-step integration process that builds upon existing integration methods of Harmony^13^ and Seurat^14^. Harmony and Seurat are widely used and well-established integration methods, known for their computational efficiency. Consequently, Smmit requires minimal effort for implementation and benefits from the computational efficiency of these two methods. Smmit can be applied to various types of single-cell multi-omics data, including datasets with joint profiling of gene expression and protein abundances (e.g., from CITE-seq^1^) as well as joint profiling of gene expression and chromatin accessibility (e.g., from 10x Multiome or SHARE-seq^2^). Interestingly, in real data analyses, we found that the simple approach employed by Smmit often leads to better integration results compared to existing methods. These findings suggest that Smmit is an effective, generalizable, and scalable method for integrating large-scale single-cell multi-omics data.

## Results

### Smmit overview

Smmit takes as input single-cell multi-omics data from multiple samples. Smmit first employs Harmony to integrate multiple samples within each data modality. We selected Harmony as the integration method due to its strong performance reported in recent benchmarking studies^15,16^, its scalability, and its compatibility with the Seurat pipeline. The Harmony integration step removes unwanted sample-specific effects while preserving cell type differences.

The Harmony-integrated reduced-dimension representations are then fed into Seurat’s weighted nearest neighbor (WNN) function to integrate multiple data modalities and produce a single UMAP. Smmit returns a unified Seurat object containing the WNN outputs, which can be directly used in the standard Seurat pipeline for downstream analyses, such as cell clustering, cell type identification^17^, and differential analysis.

### Integration of a single-cell Multiome dataset

We applied Smmit, along with five competing methods (scVAEIT, Multigrate, scMoMaT, MultiVI, and MOFA+), to a benchmark dataset generated using the 10x Single-Cell Multiome platform (Methods). The dataset consists of 69,249 bone marrow mononuclear cells (BMMCs) from 10 healthy human donors. Cell type annotations provided in the original dataset were treated as the gold standard. Each sample was considered a separate batch.

We obtained low-dimensional representations learned by each method after integrating data across samples and modalities. Cell clustering was then performed on the integrated low-dimensional representations. Figures 1a and 1b display cells from different samples and annotated cell types in UMAP space for each method. Figure 1c shows kBET (Methods), a quantitative metric evaluating the extent to which batch and samples effects were removed during the integration process. Figure 1d shows the adjusted Rand index (ARI, Methods), another quantitative metric assessing the consistency between computationally derived cell clusters and annotated cell types.

**Figure 1.**
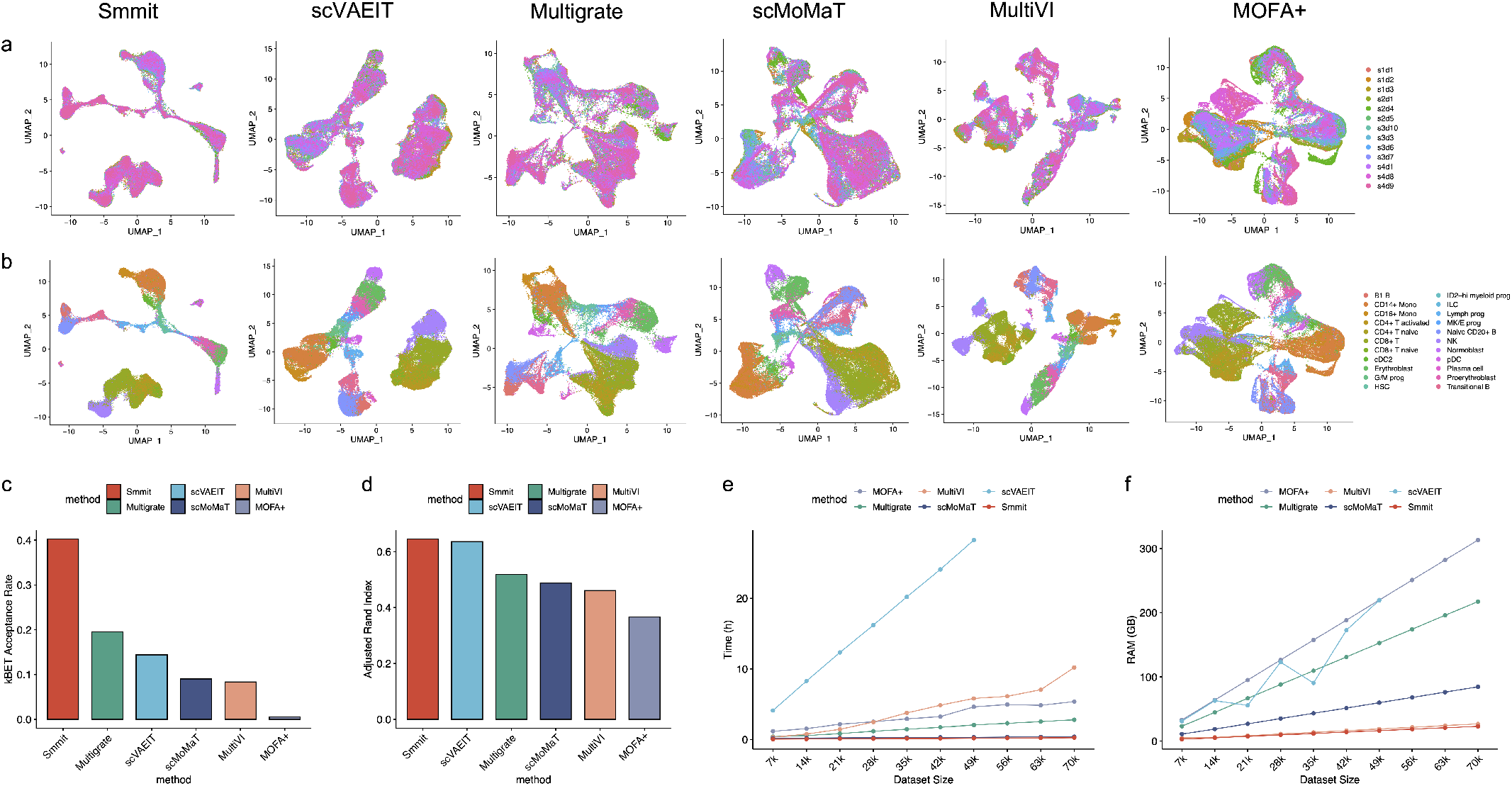
Integration of a single-cell Multiome dataset. **a-b**, UMAP plots of integrated low-dimensional representations generated by different methods, colored by individual samples (**a**) and annotated cell types (**b**). **c**, kBET acceptance rate for each method. **d**, ARI score for each method. **e**, Running time for each method (y-axis) across varying numbers of cells (x-axis). **f**, Memory usage for each method (y-axis) across varying numbers of cells (x-axis).

Smmit demonstrates the best integration results compared to other methods. While other methods generate clusters of cells that are confined to one or a small number of samples, Smmit effectively mixes cells from different samples (Figure 1a), as evidenced by its highest kBET metric among all methods (Figure 1c). Additionally, Smmit accurately assigns cells of the same cell type to the same position in UMAP space (Figure 1b), achieving the highest ARI (Figure 1d). In contrast, other methods often assign cells of the same cell type to multiple, separated positions in UMAP space (Figure 1b). The UMAP pattern also reflects the overall process of hematopoietic stem cell (HSC) differentiation into lymphoid, myeloid, and erythroid cells. These results demonstrate that Smmit excels in removing technical effects across batches while preserving the biological signals in the data.

We also compared the computational efficiency of these methods (Methods), including their running time (Figure 1e) and memory usage (Figure 1f). Smmit exhibited the shortest running time and the smallest memory usage among all methods. It completed the computation within 15 minutes and required only 23.05 GB of memory to process 70,000 cells. In contrast, Multigrate and scVAEIT, which rank among the top three best-performing methods, required 2.79 hours and over 28.26 hours of running time, and 217.29 GB and more than 230 GB of memory, respectively. These results highlight the computational efficiency of Smmit when handling large datasets.

### Integration of a CITE-seq dataset

We further applied Smmit and three competing methods (Multigrate, scVAEIT, and totalVI) to another benchmark dataset generated using the CITE-seq platform (Methods). This dataset consists of 78,651 BMMCs from 10 healthy human donors. As with the previous dataset, we evaluated the performance of each method using the cell type annotations from the original dataset as the gold standard. Each sample was considered a separate batch.

Smmit again demonstrates the best integration results. It effectively mixes cells from different samples (Figure 2a) while preserving cells of the same type in consistent positions in UMAP space (Figure 2b), leading to the highest kBET (Figure 2c) and ARI metrics (Figure 2d). In contrast, UMAP spaces generated by other methods show cells from different samples as separated, with cells of the same type assigned to different locations, resulting in inferior integration performance. Additionally, Smmit is among the most computationally efficient methods, as indicated by its running time (Figure 2e) and memory usage (Figure 2f). These results suggest that Smmit remains the best method on the CITE-seq dataset.

**Figure 2.**
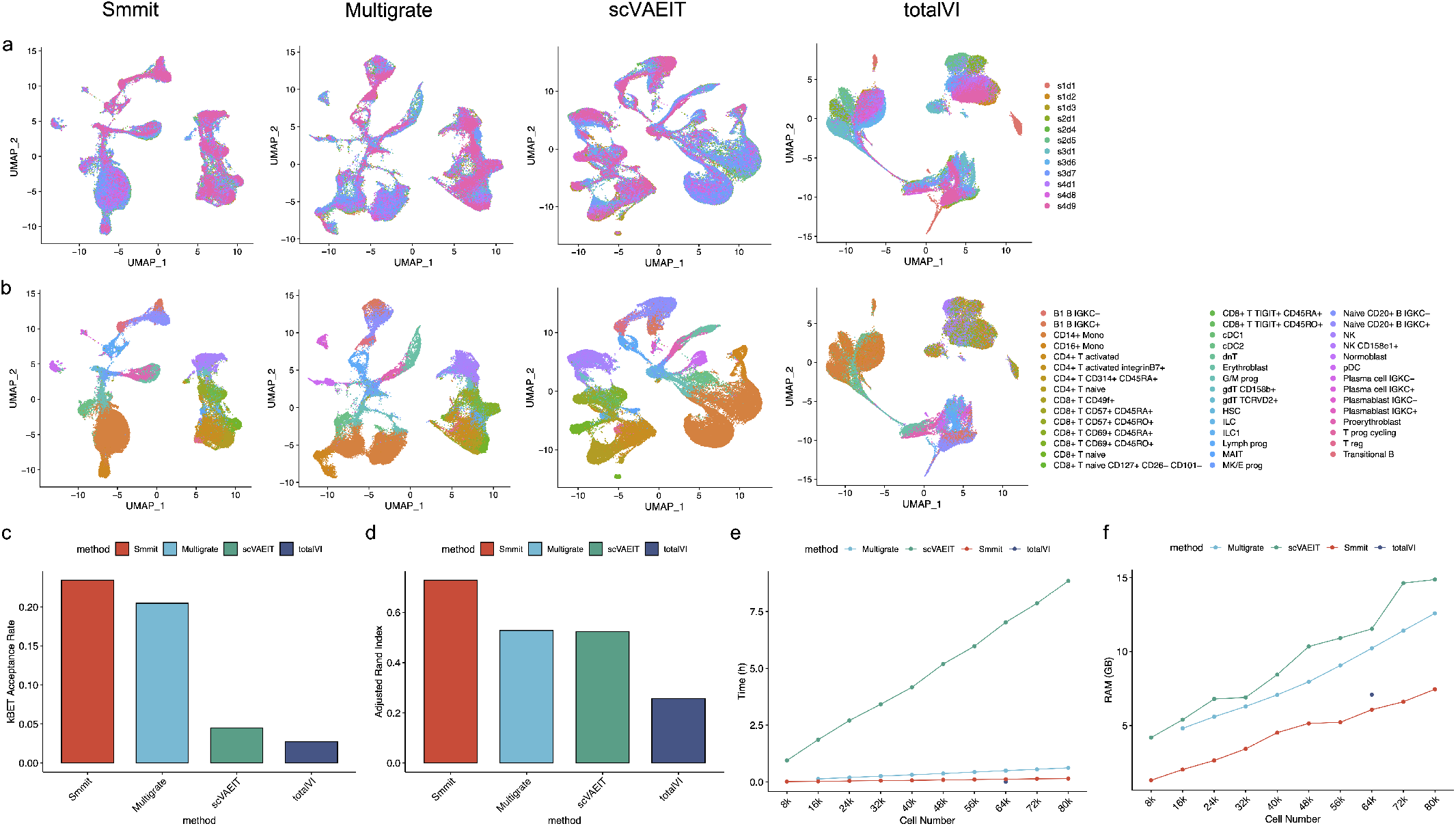
Integration of a CITE-seq dataset. **a-b**, UMAP plots of integrated low-dimensional representations generated by different methods, colored by individual samples (**a**) and annotated cell types (**b**). **c**, kBET acceptance rate for each method. **d**, ARI score for each method. **e**, Running time for each method (y-axis) across varying numbers of cells (x-axis). **f**, Memory usage for each method (y-axis) across varying numbers of cells (x-axis).

## Conclusions

We developed Smmit, a method for integrating single-cell multi-omics data from multiple samples. We demonstrated that Smmit produces superior integration results in real data applications. Smmit builds on existing computational methods, does not require GPUs, and is highly efficient for handling large datasets. These features make Smmit a user-friendly option for integrating data from multiple single-cell multi-omics samples.

## Methods

### Smmit pipeline

The input for Smmit is a list of Seurat objects prepared by the user. Each Seurat object contains single-cell multi-omics data from a single sample, processed by the standard Seurat^14^ or Signac^18^ pipeline. The Seurat or Signac objects only need to include the count data. No downstream processing, such as data normalization and scaling, is required, as this will be handled internally within Smmit. Smmit then uses the `Seurat::merge()` function to merge the input list of Seurat objects into a single merged Seurat object. For antibody-derived tags (ADT) in CITE-seq^1^, only proteins that are shared across all Seurat objects are used in the merging process.

For merged RNA data, the following Seurat functions are applied sequentially with default parameters to produce a joint principal component (PC) space: `NormalizeData(), FindVariableFeatures(), ScaleData(), RunPCA()`. For merged ADT data, the following Seurat functions are applied sequentially to produce a joint PC space: `NormalizeData(), ScaleData(), RunPCA()`. The `NormalizeData()` function is executed using a centered log ratio transformation that normalizes across cells, while the other two functions employ default parameters. For merged ATAC data, the following Signac functions are applied sequentially with default parameters to produce a joint latent semantic indexing (LSI) space: `FindTopFeatures(), RunTFIDF(), RunSVD()`. Within each modality, the `harmony::RunHarmony()` function is then applied to the joint space of reduced dimensions to integrate samples.

Finally, the Seurat function `FindMultiModalNeighbors()` is used to integrate the Harmony integrated reduced dimensions of RNA and ADT or RNA and ATAC. For RNA and ADT integration, if ADT has less than 30 PCs, all PCs in ADT are used. Otherwise, top 30 PCs in ADT are used. The number of PCs used for RNA and ADT is set to be the same. For RNA and ATAC integration, the top 30 PCs are used for RNA, and the top 2 to 30 LSI components are used for ATAC. The Seurat function `RunUMAP()` is then used to produce a UMAP space based on the WNN results.

### Data processing

To demonstrate the utility of Smmit, a processed 10X Multiome dataset containing 13 samples from 10 donors of BMMCs was downloaded from^19^. Cells with at least 60 ATAC read counts and at least 100 RNA read counts were retained. Additionally, a processed CITE-seq dataset with 12 samples from 10 donors was downloaded from the same study. Cells with positive RNA expression in at least 100 genes were retained.

### Competing methods

The following methods were used to analyze the Multiome data: scVAEIT (version 0.0.0), Multigrate (included in scArches version 0.6.1), scMoMaT (version 0.2.2), MultiVI (included in scvi-tools version 0.19.0), and MOFA+ (version 0.7.1).

The following methods were used to analyze the CITE-seq data: Multigrate (included in scArches version 0.6.1), scVAEIT (version 0.0.0), and totalVI (included in scvi-tools version 0.19.0).

We followed the settings and parameter choices described in a previously published benchmark paper to implement these methods^20^.

### Evaluations

#### kBET acceptance rate

The k-nearest-neighbor Batch Effect Test (kBET) is used to evaluate how effectively the batch effect has been removed during batch correction. It quantitatively measures whether the composition of batch labels among the k-nearest neighbors of a cell matches the overall batch label distribution in the dataset. kBET values are computed using the scIB package (version 1.1.5) with a nearest-neighbor graph as input.

For Smmit, we first computed the modality weights of Harmony embeddings from RNA and ATAC/ADT individually using the function `Seurat:::FindModalityWeights`, with all other parameters set to their default values. The final nearest neighbor graph was computed using the function `Seurat:::MultiModalNN`. Since all other methods provide lower-dimensional embeddings, we used the `Seurat::FindNeighbors` function to directly generate the neighborhood graph.

#### Adjusted rand index

Rand index (RI) is defined as

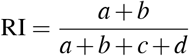

where a is the number of pairs of points that are in the same cluster in both clusterings, b is the number of pairs of points that are in different clusters in both clusterings, c is the number of pairs of points that are in the same cluster in one clustering but in different clusters in the other, and d is the number of pairs of points that are in different clusters in one clustering but in the same cluster in the other.

Adjust rand index (ARI) is defined as:

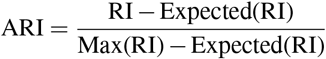

where **Expected(RI)** is the expected value of RI under random chance, and **Max(RI)** is the maximum value the Rand Index can achieve.

To compute the adjusted Rand index, we first cluster cells using the weighted nearest neighbor graph from Smmit and KNN graphs from individual methods with the `Seurat::FindClusters` function. We set a gradient of resolution ranging from 0.1 to 2 in increments of 0.1. Each algorithm is evaluated using the highest adjusted Rand index achieved under different clustering resolutions. ARI was computed using the function `fossil::adj.rand.index`.

#### Running time and memory usage

Due to their neural network-based nature, Multigrate, scVAEIT, scMoMaT, MultiVI, and totalVI were evaluated on a SYS-120GQ-TNRT GPU server with a maximum RAM of 230 GB and an Nvidia RTX A6000 GPU. Both Smmit and MOFA+ were evaluated on a UCSB-B200-M5 CPU server with a maximum RAM of 730 GB. To evaluate the time spent and RAM allocated on datasets of different sizes, we sampled cells both Multiome and CITE-seq datasets from 10% to 90% in increments of 10% based on the batch label in each dataset. We then recorded the user time and maximum resident set size allocated during the execution of each method.

## Acknowledgments

This study was supported by the National Institutes of Health under Award Number U54AG075936 and R35GM154865.

## Author contributions

C.W. and Z.J. conceived the study. C.W. conducted the analysis and developed the software. C.W. and Z.J. wrote the manuscript.

## Data availability

Both the processed 10X Multiome and CITE-seq datasets were downloaded from the Gene Expression Omnibus (GEO) under the accession GSE194122.

